# Impact of *Babesia microti* on the initiation and course of pregnancy in murine model of vertically transmitted infection

**DOI:** 10.1101/2020.10.12.335927

**Authors:** Katarzyna Tołkacz, Anna Rodo, Agnieszka Wdowiarska, Anna Bajer, Małgorzata Bednarska

## Abstract

Genus *Babesia* groups tick-transmitted protozoa causing babesiosis, a malaria-like disease. Vertical transmission of *Babesia* spp. was reported in mammals, however, the exact timing and mechanisms involved in this mode of transmission are not currently known. In this experimental study we evaluated: 1) the reproductive success, and success of vertical transmission of *Babesia microti* in mice mated in acute and chronic phases of the infection and in pregnant mice infected during early and advanced pregnancy; 2) possible influence of the pregnancy on the course of parasite infection (parasitaemia) in females; and 3) pathological changes in females and their embryos induced by infection. Blood smears and PCR targeting the 550 bp 18S rRNA gene fragment were used for the detection of *B. microti*. Histopathological examination was performed on collected tissues.

Successful development of pregnancy was recorded only in females in the chronic phase of infection. The success of vertical transmission of *B. microti* in this group was 63% (71/112). In females mated in the acute phase of infection or on the 4^th^ day of pregnancy, no evidence for pregnancy development were observed. In the group infected on the 12^th^ day of pregnancy, numerous complications including pregnancy loss and stillbirth were recorded. During the acute phase of infection, parasitaemia was lower in pregnant females in comparison to infected, non-pregnant control females.

Acute *B. microti* infection prevents pregnancy initiation and development of pregnancy at a very early stage, and causes severe complication in BALB/c mice in the second and third trimester of pregnancy. Chronic *B. microti* infection has no negative impact on the initiation and development of pregnancy, but resulted with congenital infections. Further study is required to determine to what extent maternal antibabesial immune responses and potential placental accumulation of parasites contribute to compromised pregnancy in the murine model of congenital *Babesia* infection.

**Author summary:** The mouse is the most common mammalian model for studying human parasitic diseases, including malaria, toxoplasmosis, Chagas disease, and babesiosis. Babesiosis is an emerging intraerythrocytic infection caused by protozoal parasites, mostly *Babesia microti*. Our previous work in murine model proved that vertical transmission of *Babesia microti*, is a third way - after tick-bite and blood/organ transfusion - to acquire babesiosis. In this study we focused on investigating how the infection influences the course of pregnancy. We were interested in how variations in infection acquisition time and infection phase influence the reproductive success of mice and vertical transmission of parasites. We expected that the infection causes severe pathological changes in the organs of infected females and their offspring. Results obtained in this study have shown that vertical transmission of *B. microti* was only possible in chronically infected mice, in which health status and reproductive success were not compromised by the infection. Acute infection made successful reproduction impossible, however, the infection had no significant effect on the histopathological condition of tissues. We hope that these insights into *B. microti* vertical transmission will lead to the better understanding of congenital babesiosis.

## Introduction

Parasitic infection may have negative effect on host breeding success and offspring’s health condition causing congenital infections. Among the most well-recognised congenital parasite infections are those of *Toxocara canis* in dogs, and *Toxoplasma gondii, Trypanosoma cruzi*, and *Plasmodium* spp. infections in humans [1–4].

Genus *Babesia* groups tick-transmitted protozoa responsible for babesiosis, a malaria-like disease in humans and animals [5]. Acute babesiosis can lead to death, nevertheless, asymptomatic infections are also reported from humans and animals [6, 7]. Asymptomatic infection in females may last for years and have negative impact on breeding, resulting with congenital infection. Congenitally acquired babesiosis was reported for several *Babesia* species. Inborn babesiosis was recognised in livestock [8–11] and dogs [12, 13]. Congenital infection was also reported in small rodents known as the reservoir hosts of these parasites [14, 15]. Experimental studies supported vertical route of transmission for *B*. *gibsoni* in dogs [16] and *B*. *microti* in mice [17].

Reported cases of inborn babesiosis differ in their outcome. Congenital infections in voles *Microtus* spp. and *Peromyscus leucopus* were asymptomatic [14, 15], while infection in dogs resulted with stillbirth or the development of fully-symptomatic disease [12, 13, 16]. A newborn calf infected with *B. bovis* died without treatment in 24 hours after delivery [8].

In our well established experimental model of vertical babesiosis in mice, the course of *B. microti* infection in the offspring was highly dependent on the phase of the infection in the females [17]. Females with chronic infection gave birth to congenitally infected pups with asymptomatic infection, while those in the acute phase did not deliver any offspring [17]. In humans, currently six cases of congenital babesiosis due to *B. microti* have been reported [18–24]. In all cases, the symptoms of the disease in children occurred several weeks after delivery. Infants presented flu-like symptoms, anaemia, and hepatomegaly [18–24]. In order to prevent and effectively treat congenital babesiosis, it is necessary to recognise the mechanisms of vertical transmission of piroplasm (i.e. exact place, time, pathogenesis) and the consequences of parasitic infection at different phases of pregnancy.

The aims of this study are to: 1) evaluate and compare the success of vertical transmission of *B*. *microti* in mice mated in acute and chronic phases of infection, and in pregnant mice infected during early and advanced pregnancy; 2) evaluate and compare reproductive success of mice infected before and during pregnancy; 3) compare the course of parasite infection (parasitaemia) in female mice infected before and during pregnancy; and 4) assess and compare pathological changes in female mice and their embryos induced by parasite infection.

Results obtained in this study have shown that vertical transmission of *B. microti* was only possible in chronically infected mice, in which health status and reproductive success were not compromised by the infection. Acute infection made successful reproduction impossible, however, the infection had no significant effect on the histopathological condition of tissues.

## Materials and methods

### Animals

BALB/c mice, 82 females (experimental and control groups) and 33 males (mating), in the age 10-12 weeks were used in the course of the experiments (Table 1). BALB/c mice are medium-resistant to *B. microti* infection, which has made them a suitable model for human babesiosis studies for many years [25, 26]. The parasite can be maintained in adult BALB/c females for the long chronic period following recovery from acute babesiosis [7]. This strain of mice was successfully used in our preliminary studies on vertical transmission of *B. microti* [17]. All animals were kept in standard cages provided with sawdust, nest material, food, and water *ad libitum*. Females were kept in cages in groups of 2-8 before mating. Pregnant females and dams with pups were housed in pairs. It was previously demonstrated, that two females per cage raised the higher number of weanlings per cage [27]. The sire males of BALB/c strain were housed individually while not paired with females. In order to mate, males were joined with females for one night (1 male + 1-2 females). All of the procedures conducted on mice were approved by the First Ethics Committee for Animal Experimentation in Poland (ethical license numbers: 406/2013, 716/2015 and 536/2018), according to the principles governing experimental conditions and care of laboratory animals required by the European Union and the Polish Law on Animal Protection.

**Table 1.**
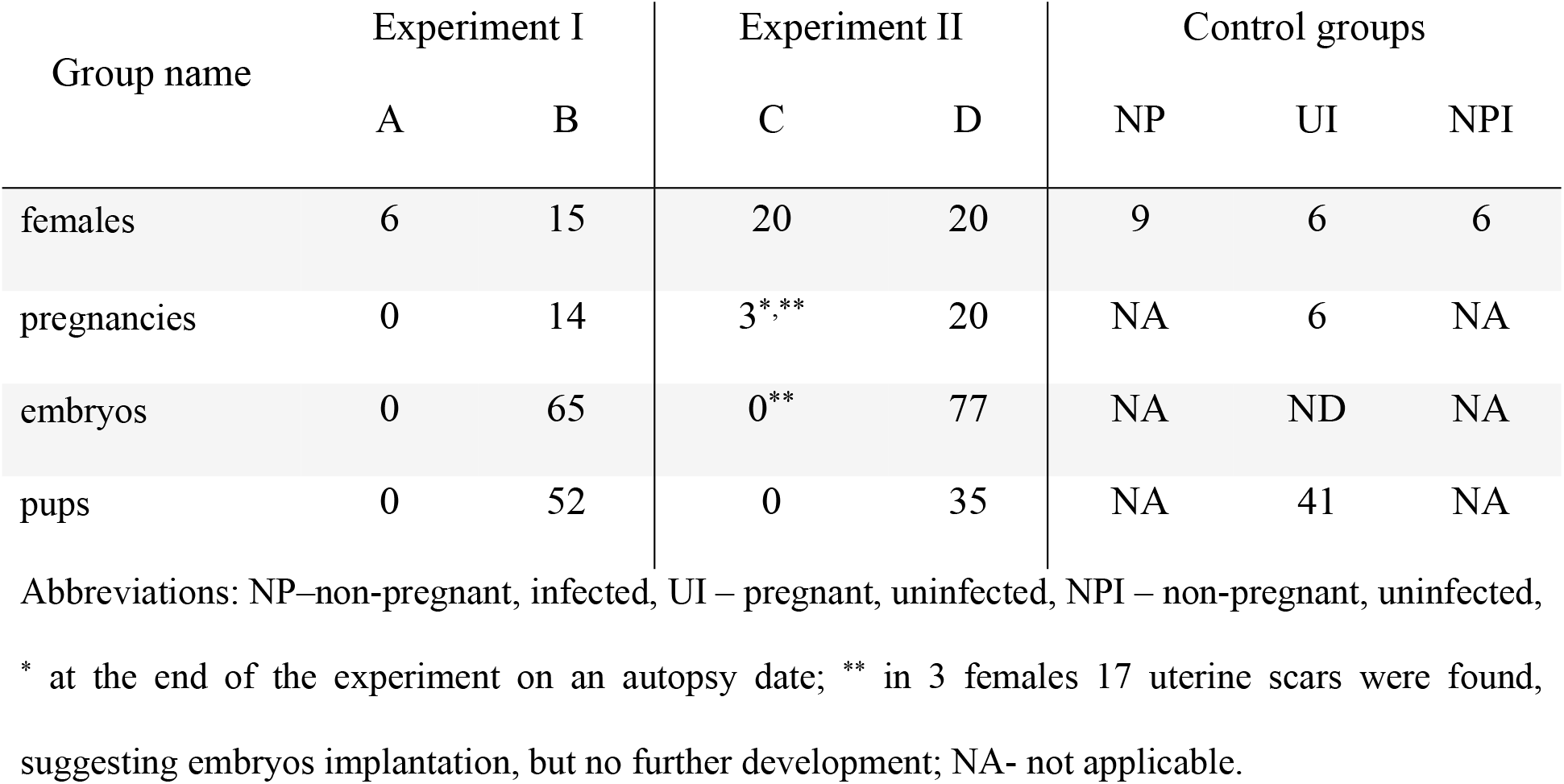
The total number of females in the experimental and control groups, and the number of embryos and pups recorded in the course of the experiments.

### *Babesia microti* strain

The female mice were infected with the *B. microti* King’s 67 strain, which originated from field voles in the Oxford area, United Kingdom [28]. The blood is passaged from infected to naive mice by intraperitoneal injections [7, 17]. Mice are infected with 5 × 10^6^ infected red blood cells (iRBCs) in the volume of 0.2 ml. This method has been successfully used in experimental studies on *Babesia* and other blood parasites worldwide [7, 17, 29, 30]

### Mating and breeding of mice

Determination of the phase of oestrous cycle in females was performed with the crystal violet staining of vaginal smears following the protocol of [31]. Females were joined with males to mate for one night. The presence of a vaginal plug the next morning indicated that insemination had occurred. Around 8-12 days post fertilisation, females were weighed and vaginal smears were performed to evaluate pregnancy development. Females were housed in pairs during pregnancy. Pups from the chronic phase of infection (Group B, see below: study design) were housed with their mothers until the end of experiment.

### Study design

The first set of experiments involved females infected with *B. microti* before mating. Females were assigned to two experimental groups:

In Group A, six females in the appropriate phase of the reproductive cycle were joined with males on the 7^th^ day post infection with *B. microti* (acute phase).

In Group B, 15 females were joined with males in the chronic phase of the *B. microti* infection after the 40^th^ day post infection.

A second set of experiments involved female mice infected with *B. microti* during pregnancy: In Group C, 20 fertilised females were infected with *B. microti* on the 4^th^ day post mating, in the early pregnancy.

In Group D, 20 fertilised females were infected with *B. microti* on the 12^th^ day post mating, in advanced pregnancy.

Three control groups of female mice were established. In total, 21 control mice were used in the experiments:

Control Group 1 (NP) of *B. microti* infection: nine virgin females were infected with *B. microti,* to monitor the course of infection in non-pregnant females (infected, non-pregnant). Control Group 2 of pregnancy (UI): six fertilised uninfected females were used to monitor blood parameters and the pregnancy development in healthy females (uninfected, pregnant). Control Group 3 (NPI): six non-pregnant, uninfected females.

### Course of the experiments

Blood from the tip of the tail was collected from each infected female to perform blood smears, starting on the day of infection until the day of the autopsy. From the 1^st^ to 20^th^ day post infection, blood was collected every 2-4 days, after which samples were collected every 10 days.

At the end of the experiments females were euthanised by cervical dislocation. Females from Group A were autopsied on the 12^th^ day of the expected pregnancy. As their weight did not change since fertilisation, and USG examination was unable to detect any developing embryos, females from this group were autopsied to check the development of pregnancy. Females from Group B were autopsied on the 12^th^ (1 female), 16^th^ (3 females), and 18^th^ (3 females) day of pregnancy, and after the delivery – on the 1^st^ (3 females), 7^th^ (2 females) and 14^th^ (3 females) day postpartum.

Females from Group C were autopsied on the 8^th^ (6 females), 12^th^ (6 females), 14^th^ (6 females), and 18-20^th^ (2 females) day of pregnancy.

Females from Group D were autopsied between the 14^th^ and 18^th^ day of pregnancy (2 females on 14^th^, 6 females on 16^th^, 6 females on 18^th^), and 6 females on the 1^st^ day postpartum.

Females from Control Group 1 (NP) were autopsied on the 15^th^ (6 females) day post infection to collect samples for morphology, histopathology, and parasitaemia evaluation. Another three mice, infected with *B. microti* to compare the course of parasitaemia with females from Group B, were autopsied at the age of 50 weeks (150 day post infection).

Females from Control Group 2 (UI, pregnant, uninfected) were autopsied on the 1^st^ (1 female), 7^th^ (3 females), and 14^th^ day postpartum (2 females).

Females from Control Group 3 (NPI, non-pregnant, uninfected) were autopsied at the age of 15 weeks - around the same age as females from experimental Groups A, C, and D (6 females).

At the autopsy of pregnant females, data on embryo development and tissue samples were collected for a range of laboratory investigations. Embryos (if present) were first isolated from the uterus, washed in sterile water, counted and weighted individually, and visually evaluated for the presence of developmental abnormalities (i.e. malformation of limbs, evidence of stillbirth/abortions).

Reproductive success was calculated as the mean number of well-developed, normal embryos/pups per female in the group. Survival rate was described as the percentage of alive embryos/pups from the total number of offspring in the litter at the day of the autopsy. Mean litter size was calculated for each group as the mean number of collected pups and embryos per female.

Females and offspring were autopsied. Blood samples (1 ml) were collected with a sterile syringe directly from the heart following the death of the pregnant females. Blood smears were prepared immediately for evaluation of parasitaemia, 100-200 μl of blood was kept frozen for DNA extraction, and remaining blood sample was used for the determination of blood parameters (selected counts). Selected females’ tissue samples were collected for evaluation of pathological changes during babesiosis and for the determination of the effect of parasite infection on the course of pregnancy. Organs (brain, heart, lungs, kidneys, liver, spleen, uterus) were isolated and weighted. A fragment of each organ was fixed in 10% formalin for histopathological examination and another fragment was frozen at a temperature of −20°C for DNA extraction.

At the autopsy of offspring, selected tissue samples were collected for identification of congenital *Babesia* infection. At each age we were able to collect heart and lung samples. Organs were isolated and weighted, then a fragment of each organ was frozen at a temperature of −20°C for DNA extraction. If possible, a fragment of organ was fixed in 10% formalin for histopathological examination

### Laboratory methods

#### Monitoring of pregnancy development

During pregnancy females were weighted every 2-4 days. Between the 12^th^ and 18^th^ day of pregnancy, ultrasound examinations were performed by an experienced veterinarian practitioner to monitor pregnancy development status in Groups B (*n* = 11), D (*n* = 10), and UI (*n* = 4), which were the only groups where we recorded advanced pregnancy. The examination was performed with an Esaote ultrasound, MayLab One, with a 18 MHz line probe at a test depth of 3-4 cm. The study was performed both in the longitudinal and transverse axis of the animal. The mice were examined without anaesthesia (to avoid any impact on the embryos), after shaving the fur on the abdomen and applying the gel for the ultrasound examination (Aquasonic). The mice were immobilised with the help of a grip on the skin of the neck. Examination was performed on the 12^th^, 14^th^, 16^th^, and 18^th^ day of pregnancy. Embryo heart rate was estimated during each monitoring session.

#### Monitoring of the course of *B. microti* infection

Parasitaemia was determined on the basis of Hemacolor® (Merck, Germany) stained blood smears as described previously [17].

#### Histopathological study and blood parameters

Histopathological specimens were analysed as paraffin sections stained with haematoxylin-eosin method for evaluation of pathological changes in tissues [32]. Blood samples (0.25-0.5 ml) were collected into the probes with EDTA. RBC and platelet counts were performed by a commercial diagnostic laboratory (LabWet, Warsaw).

#### Detection of *B. microti* infection in females (experimental) and offspring (congenital)

Genomic DNA was extracted from blood and tissue samples of females and from tissue samples of embryos using a QiagenDNeasy® Blood & Tissue Kit. Detection of *B. microti* isolates were performed by the amplification and sequencing of the 559 bp fragment of 18S rDNA by PCR (first run) and nested-PCR (in the case of no, or weak, signal from the initial one-step PCR). The primers and thermal profiles have been described previously [14, 17, 33]. Negative controls were performed in the absence of template DNA. As positive controls we used the genomic DNA of *B. canis* extracted from canine blood. PCR products were subjected to electrophoresis on a 1.5% agarose gel, stained with Midori Green stain (Nippon Genetics, GmbH, Düren, Germany).

### Statistical analyses

The adopted statistical approach has been documented comprehensively in our earlier publications [14, 17]. Analyses were carried out using SPSS v. 21.0. Student’s T-tests were used to compare: rate of *Babesia* positive pups and embryos, reproductive success, survival rate, mean litter size, mean body mass, heartbeat rates, and mean blood counts between groups.

Prevalence (percentage of animals infected) was analysed by maximum likelihood techniques based on log-linear analysis of contingency tables. For analysis of the prevalence of *Babesia* in pups, we fitted prevalence of *Babesia* infection as a binary factor (two levels: infected = 1, uninfected = 0), development stage (two levels: embryo=1, pup=2), age of pups (three levels: 1=1, 7=2, 14=3 day postpartum. The prevalence of *B. microti* is presented as percentages with 95% confidence interval (CI_95_) calculated with bespoke software based on the tables of Rohlf & Sokal, by courtesy of F. S. Gilbert and J. M. Behnke [34].

General linear model (GLM) was used for comparison of parasitaemia of *B. microti*, which is reported with standard errors of their means (SE). Mean parasitaemia of *B. microti* infection was calculated as the number of iRBCs/1000 RBCs. When samples were only positive by PCR, an intensity of 0.001 iRBC was implemented into quantitative statistical analysis.

## Results

### Reproductive success of *B. microti*-infected female mice

Female mice in Control Group UI showed no complications in pregnancy development and pup delivery. All six females developed pregnancies and altogether 41 pups were obtained with the mean litter size 6.8±1.0 (Tables 1–2). USG monitoring has shown no malformations in embryos or pregnancy termination in any of the females from this group. All pups were in good health condition following delivery until the day of autopsy (Table 2). In Experiment I, the difference in reproductive success between the experimental Groups A and B was significant (*t* = −*5.42; df*= 19, *p* < 0.001). No embryos or pups were observed in females from Group A (reproductive success = 0), mated in acute phase of *B. microti* infection. Moreover, among the six females in this group, no signs of implantation of embryos were noted during autopsy (Table 2). The development of pregnancies was recorded in females from Group B, mated in chronic phase of *B. microti* infection, and the proportion of pregnant females in Group B (14/15= 93%) was similar to the proportion of pregnant females in Control Group UI (6/6 = 100%) (Tables 1–2). Reproductive success (No. of alive offspring/female, Table 2) in this group was similar to the success in the Control Group UI (7.5±0.9 vs 6.8±1.0, respectively; NS). Females in Group B presented no evidence of complications during pregnancy, their mean body weight on the 18^th^ day of pregnancy was similar to the mean weight of the females in the Control Group UI: 34.85±4.01 g and 36.66±0.88 g, respectively (NS). Ultrasound monitoring showed normal pregnancy development; growth of embryos and their heart rates (beats/minute) were normal, and there were no significant differences in comparison to embryos of females from Control Group UI (205.96±10.38/47.44±4.84 vs 195.25±13.15/47.23±5.14 on the 18^th^ day of pregnancy, respectively). Altogether, 65 normal embryos and 52 pups (117 individuals in total) were derived from females in Group B. The mean litter size was similar in Groups B and UI (7.8±0.9 vs. 6.8±1.0, respectively; NS, Table 2). Survival rate (*n* of alive offspring/*n* of all recorded offspring) was 96% in Group B (two embryos were found resorbed in uterus of female autopsied on the 18^th^ day of pregnancy and three pups were killed by the dam between the 2-4^th^ day postpartum). Pups were in good condition, the course of congenital babesiosis was asymptomatic, no malformations were observed. The survival rate was similar to the survival of the offspring in the Control Group UI (Table 2). At the age of 14 day postpartum, mean weight of pups from infected females from Group B was similar to the mean weight of pups delivered by uninfected females from Control Group UI (6.9±0.2 g vs. 6.8±0.2 g respectively; NS).

**Table 2.**
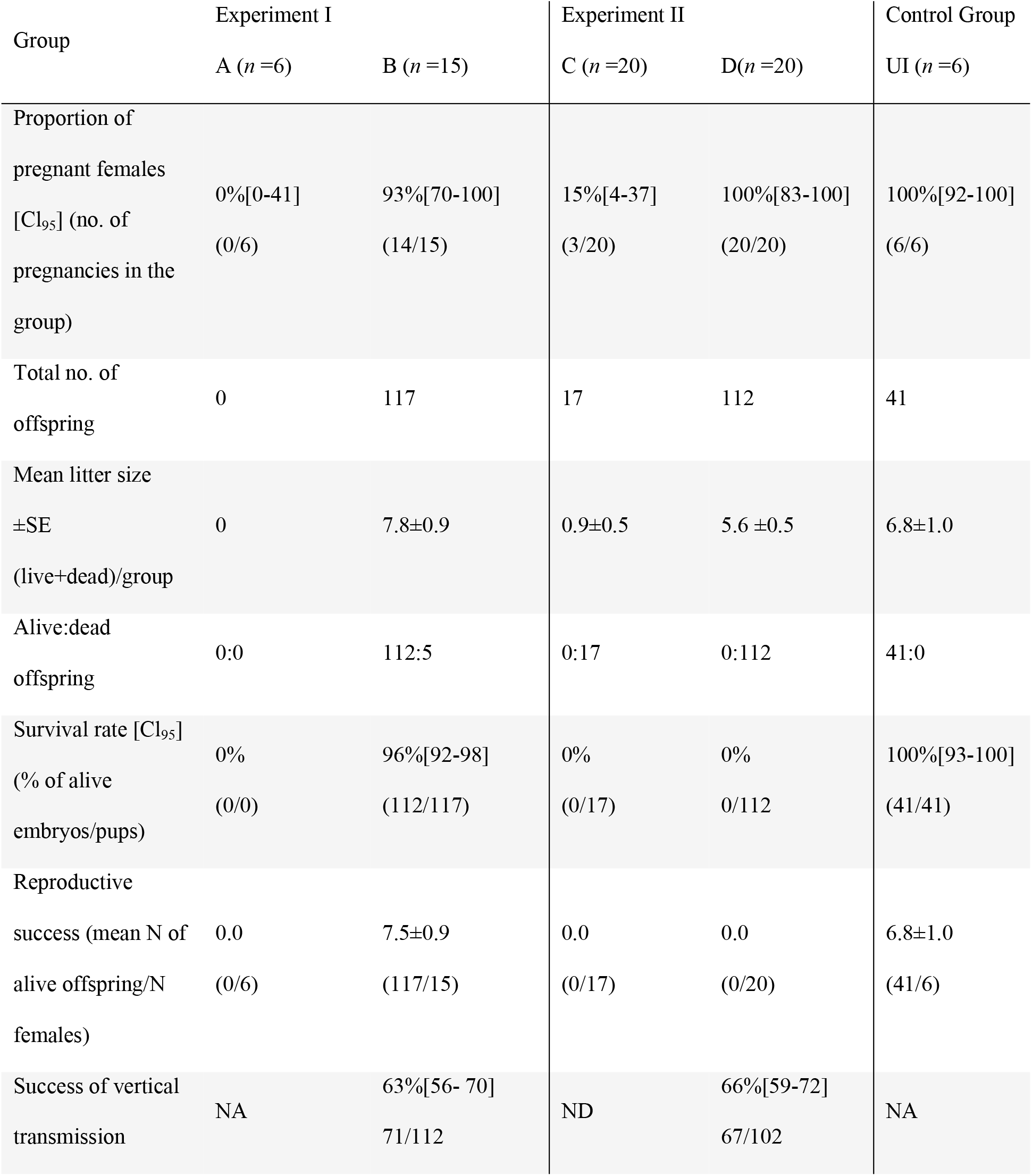
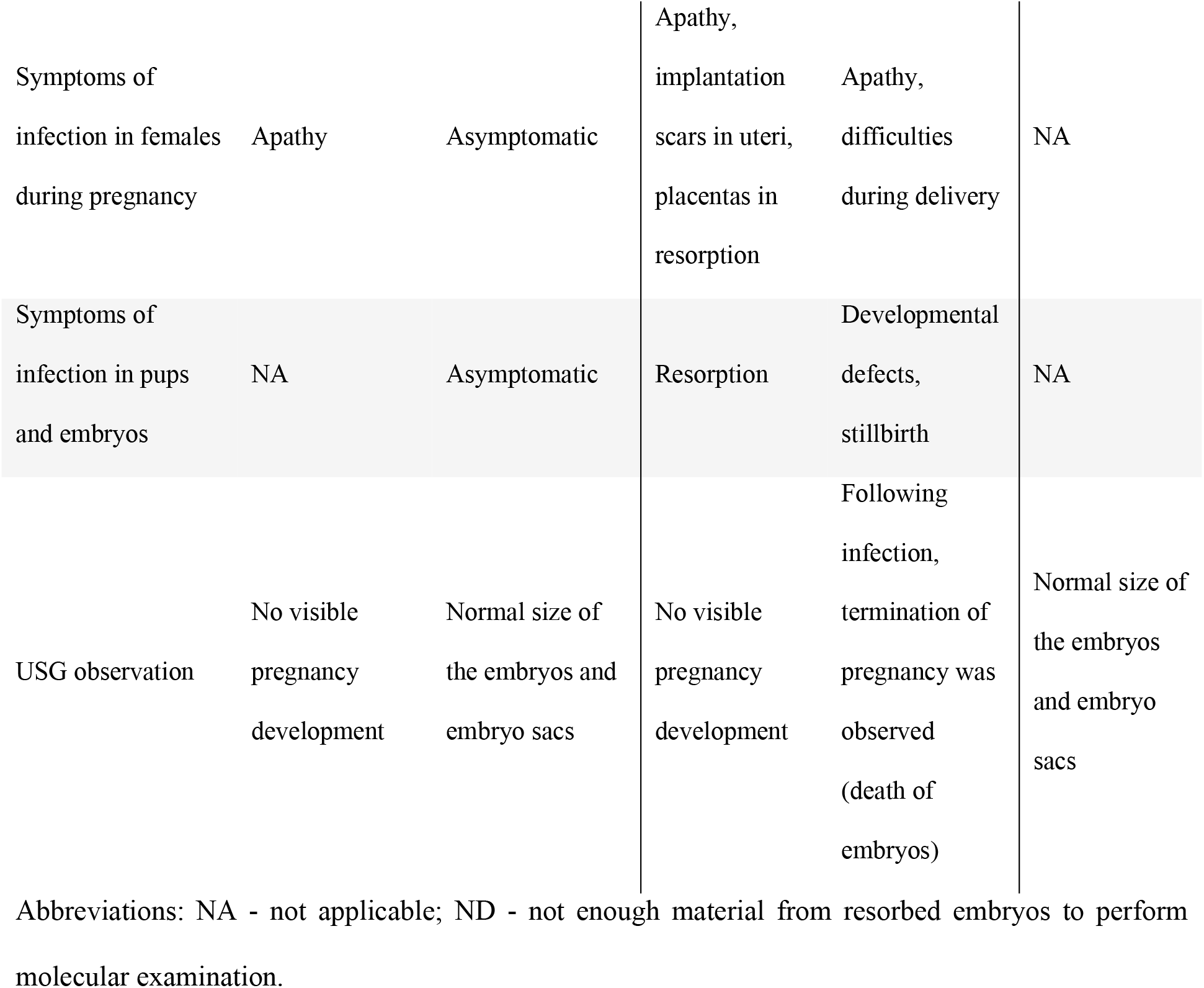
Comparison of the reproductive success of females in experimental and control groups, and features of congenital *B. microti* infection in their offspring.

In Experiment II, females were infected with *B. microti* on the 4^th^ (Group C) and 12^th^ (Group D) day of pregnancy. In Group C, infection with *B. microti* resulted in the lack of registered embryos. Only in three females from this group (15%) uterus scars were found, suggesting the implantation of 17 embryos in total. No embryos (only uterus scars) were recorded during the autopsy (undertaken between the 10^th^ and 12^th^ day of expected pregnancies). In Group D, the development of pregnancies until the 12^th^ day of pregnancy was normal (living embryos recorded during USG monitoring in all females). Heart rates of embryos during the USG examination were similar to the heart rates of embryos from the Control Group UI (204.66±7.44/39.36±4.82 and 197.04±12.71/48.98 ±4.62, respectively, NS). Females gained weight and showed no signs of complications until the 12^th^ day of pregnancy. However, numerous complications appeared after the infection with *B. microti* (Table 2). Although in Group D altogether 77 embryos and 35 pups were derived, no living offspring was obtained. USG monitoring showed that embryos that were alive on the day of infection were dying on consecutive days post infection (between the 2^nd^ and 6^th^ day post infection). The embryos’ hearts stopped beating and females had problems with delivery – pups were stillborn or died shortly post partum. As no living offspring were obtained in experimental Groups C and D, females had no reproductive success (0% of alive offspring) in comparison to the Control Group UI (100%; Table 2).

On the 1^st^ day postpartum, the mean weight of pups from Group D was half the mean weight of the pups from the Control Group UI (0.62±0.08g vs 1.31±0.13g; *t*=−4.78, *df*= 34, *p*<0.001). A number of visible malformations were recorded in embryos during autopsies in 8 females from Group D – limbs or heads were in the process of resorption.

### The success of *B. microti* vertical transmission

Embryos and pups were obtained only from females from experimental Groups B and D. The DNA of *B. microti* was detected in blood and tissue samples (collected on the 12^th^, 14^th^, 16^th^, and 18^th^ day of pregnancy, and on the 1^st^, 7^th^, and 14^th^ day postpartum) of 58% [Cl_95:_ 46-69%] (41/71) embryos and 73% [57-86%] (30/41) pups from Group B, thus the difference in prevalence of *B. microti* infection was insignificant. Overall success of vertical transmission in Group B was 63% [56-70%](Table 2). There was a significant difference in the percentage of infected individuals according to the age of pups– the older the pups were, the higher the prevalence (*Babesia* infection × day postpartum: *χ*^*2*^=9.75, *df*= 2, *p*<0.05). During autopsies performed on the 1^st^ day postpartum, 43% [21-68%] of tested pups were infected with *B. microti,* while the prevalence was higher on the 7^th^ (88% [50-99%]) and 14^th^ day postpartum (90% [68-98%], Fig 1). There was no sex difference in the frequency of congenital infection in pups −71% [46-88%] in males vs 84% [68-98%] in females (NS).

**Fig 1.**
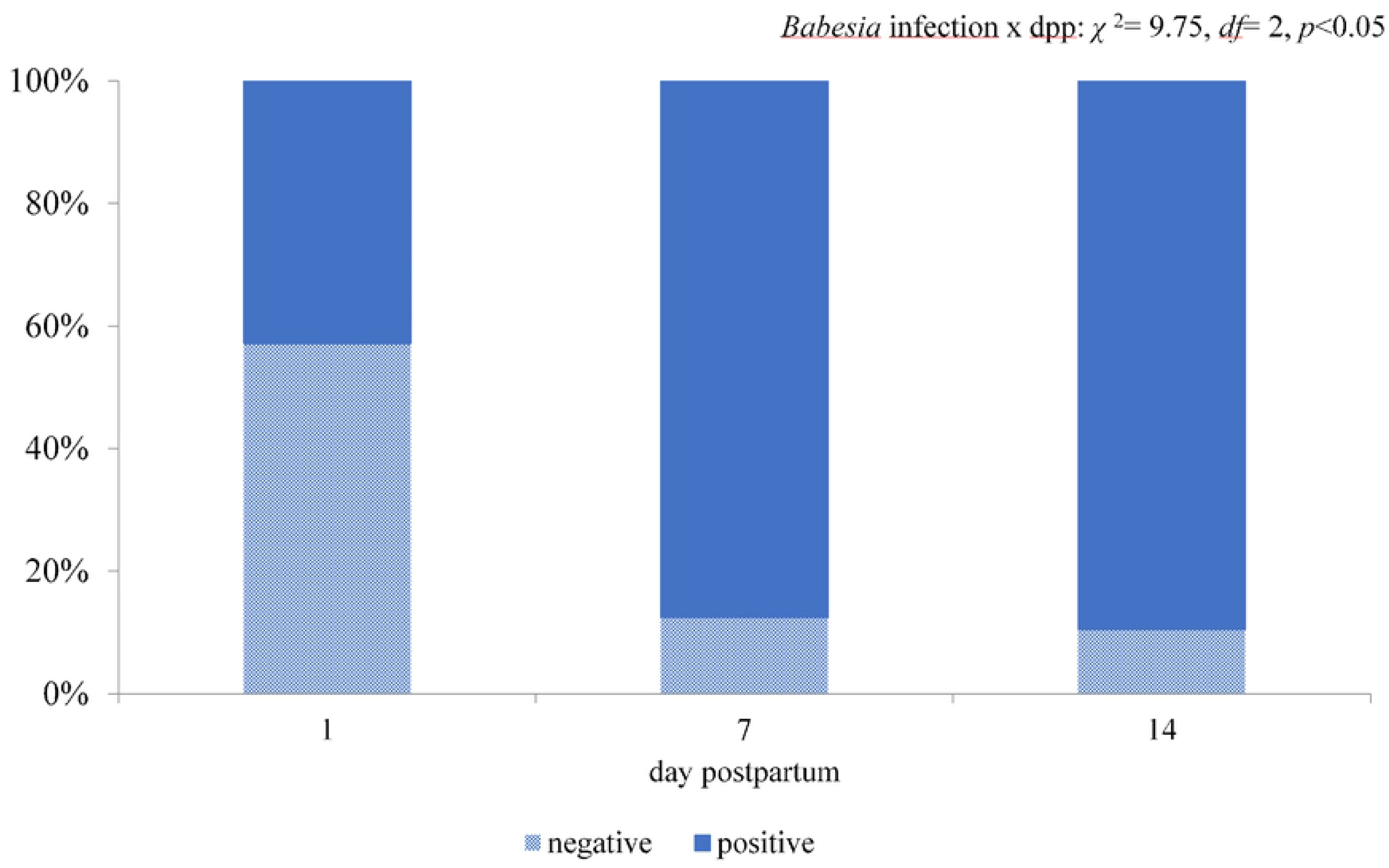
The frequency of *B. microti* infection among pups of the chronically infected females. Comparison of the frequency of *B. microti* infection among pups delivered by females in the chronic phase of infection (Group B) autopsied on the 1^st^, 7^th^, or 14^th^ day postpartum.

The frequency of detection of *B. microti* DNA in tissues of offspring and placentas obtained from experimental Groups B and D is shown in Table 3 (description further in the text). No symptoms of the acute phase of the *Babesia* infection (dark-coloured urine, febrile seizures, anorexia, apathy) were noted among the infected pups from Group B (Table 2).

**Table 3.** Comparison of the frequency of the detection of *B. microti* DNA in the tissue of offspring and placentas obtained in experimental Groups B and D.

| Speciment tested | Group B |  | Group D |  | Statistical values |
| --- | --- | --- | --- | --- | --- |
|  | % [95% CI] | infected/tested | % [95% CI] | infected/tested |  |
| Blood | 63% [56-70] | 71/112 | 66% [59-72] | 67/102 | NS |
| Brain | 15% [6-21] | 6/41 | 7% [0-30] | 1/15 | NS |
| Heart and lungs | 63% [56-70] | 71/112 | 63% [50-75] | 59/93 | NS |
| Liver | 13% [7-23] | 9/69 | 53% [29-78] | 8/15 | $t=-3.77^* df=82$ |
| Spleen | 16% [7-32] | 7/44 | 63% [29-89] | 5/8 | $t=-3.08^* df=50$ |
| Kidneys | 13% [5-29] | 6/45 | 60% [33-81] | 9/15 | $t=-4.02^* df=58$ |
| Placenta | 69% [53-82] | 29/42 | 98% [93-100] | 57/58 | $t=-4.53^{**} df=98$ |

\* $p<0.05$ t-test; \*\* $p<0.001$ t-test

In Group D, altogether 77 embryos and 35 pups were obtained. DNA of *B. microti* was detected in 66% [59-72%] of tested offspring (67 positive/102 tested): 49% [38-61%] (34 positive/69 tested) in embryos and 100% [92-100%] (33/33) in examined pups (*t*=−5.77, *df*=100, *p*<0.001, Table 3). Tissues of six embryos were too decayed to enable DNA isolation. The real success of *B. microti* vertical transmission in this group was 0% because all pups died, at latest on the 1^st^-2^nd^ day postpartum. Pups in this group presented low fitness: were stillborn, with visible head and/or limbs malformations, or died a few hours after delivery.

There was no significant difference in the frequency of vertical transmission between embryos from Groups B and D (58% [46-69%] vs 49% [38-61%], respectively, NS). In Group D, the frequency of vertical transmission was higher in the third than in the second trimester (*t*= −2.07, *df*=61, *p*<0.05, Fig 2). In the second trimester, *B. microti* DNA was detected in 13% [1-50%] of embryos and in the third trimester in 48% [38-58%] of embryos. In Group B there was no significant difference between *B. microti* DNA detection between embryos autopsied in the second and third trimesters (45% [24-68%] vs 63% [53-72%], respectively; NS). The percentage of infected pups (on the 1^st^ day postpartum) in Group D was higher than in Group B (100% vs 43%, respectively, *t*=−6.49, *df*=45, *p*<0.001)

**Fig 2.**
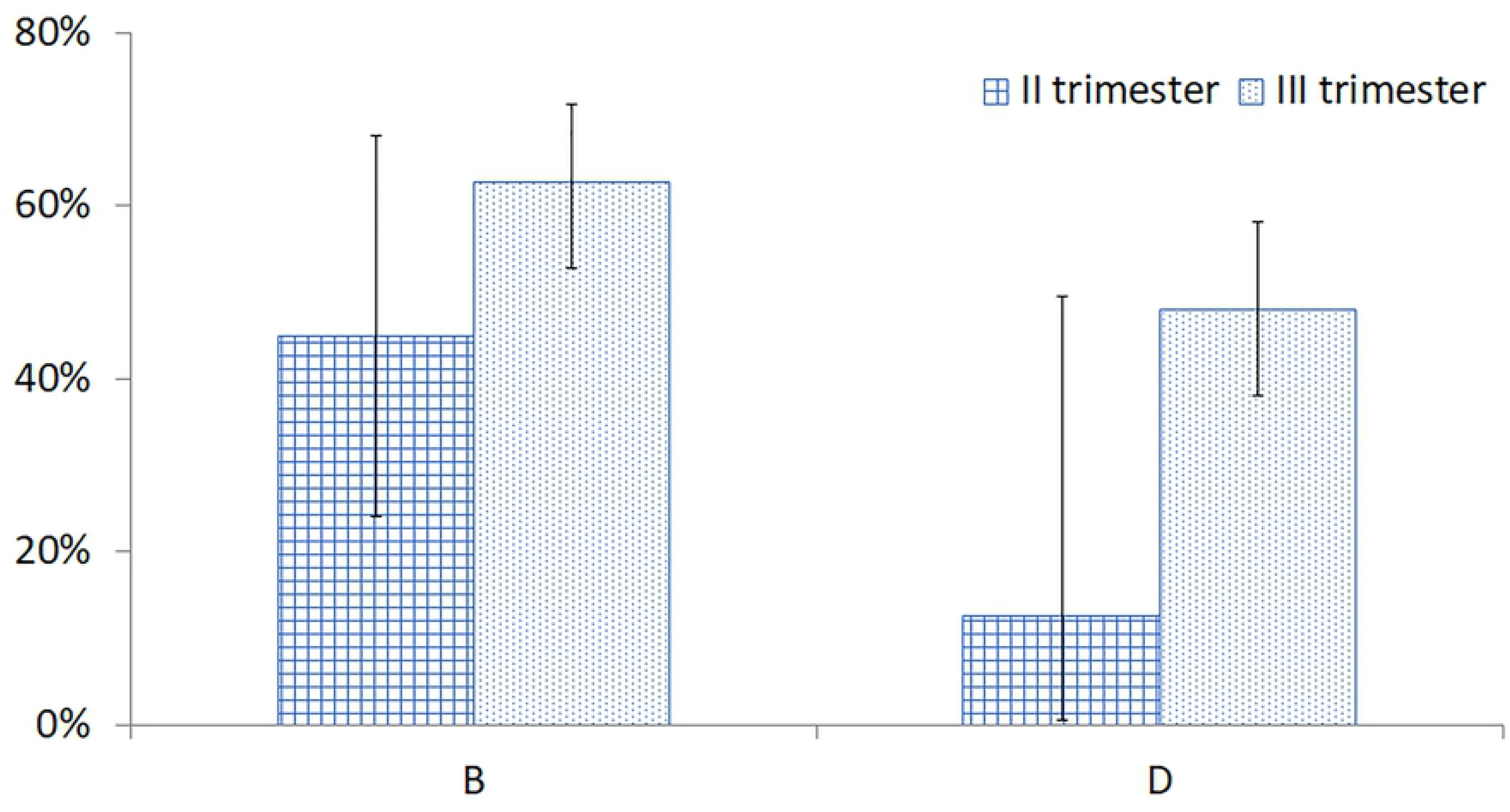
The frequency of *B. microti* infection among embryos autopsied in the second and third pregnancy trimesters. Comparison of the frequency of *B. microti* infection among embryos from Groups B and D autopsied in the second and third pregnancy trimesters.

### Comparison of the course of *B. microti* infection in females from different groups and their offspring

A comparison of the courses of *B. microti* infection during the acute and post-acute phases of infection in the experimental groups is presented in Fig 3. In all experimental groups, an acute phase of *B. microti* infection was represented by high parasitaemia during the first two weeks of infection. The highest parasitaemia was observed on the 7^th^-8^th^ day post infection in all experimental and control groups (Fig 3). Mean parasitaemia in the acute and post-acute phases of infection was the highest in the non-pregnant Control Group NP (62.94% on 7^th^ day post infection, Fig 3).

**Fig 3.**
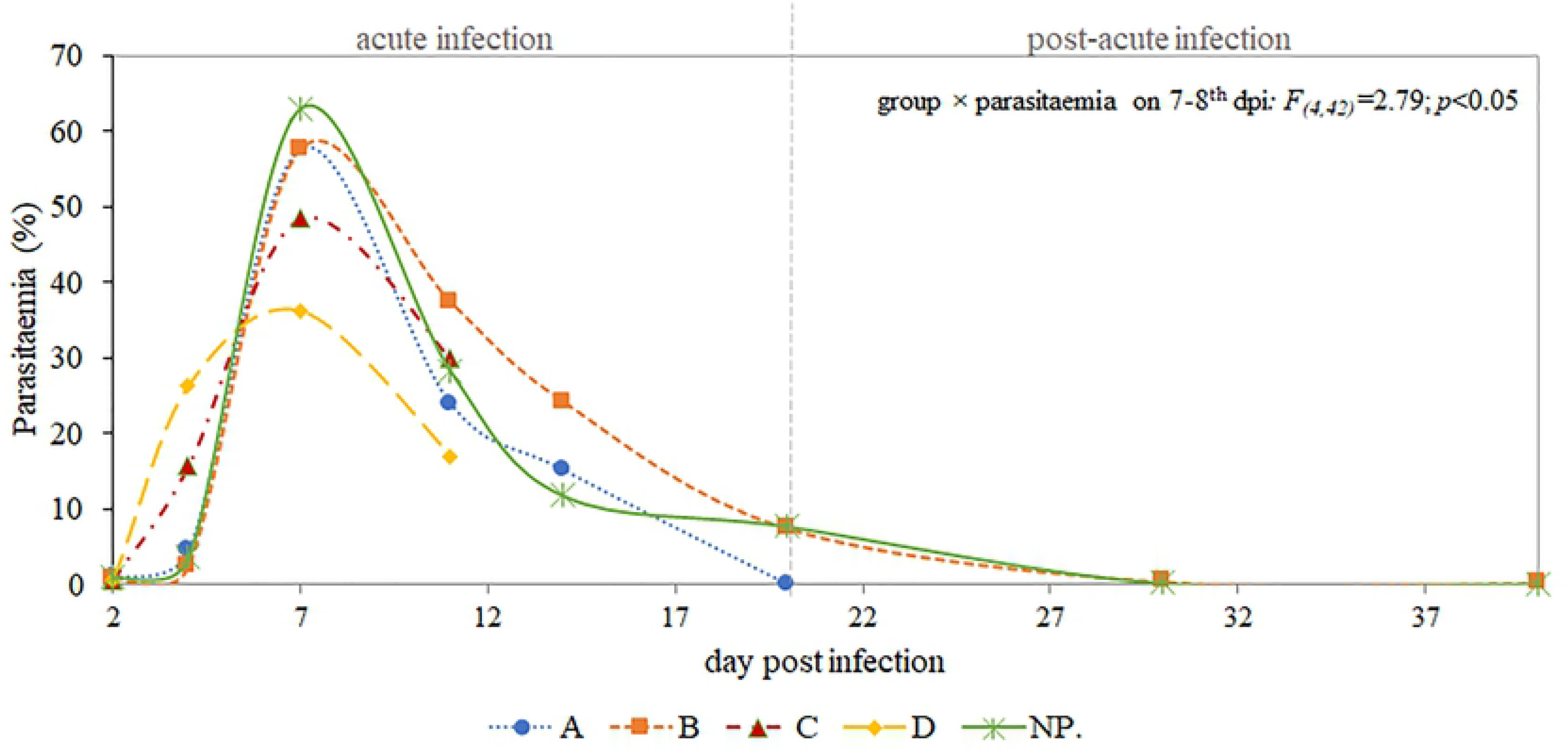
The course of *B. microti* infection in experimental and control groups. Comparison of the course of *B. microti* infection (Cl_95_) in mice pregnant in the different phases of the parasite infection. Level of *B. microti* parasitaemia on the selected day post infection. Parasitaemia is shown as a percentage of iRBCs found in mouse peripheral blood, measured on the Giemsa- stained thin smears. Datum points are the mean for three-six animals.

There was a significant difference in the mean parasitaemia of *B. microti* among experimental groups on the 7^th^-8^th^ day post infection (group ×parasitaemia on 7-8^th^ day post infection*: F_(4,42)_*=2.79; *p*<0.05, Fig 3). Maximum parasitaemia in the acute phase of infection among experimental groups was slightly higher in non-pregnant females in comparison to females with confirmed pregnancy (55.4% vs. 40.5%, respectively; pregnancy ×parasitaemia: *F_(4,42)_*=7.97, *p*<0.05). No significant associations were observed between the parasitaemia and success of vertical transmission or between parasitaemia and reproductive success of a female across the experimental groups.

### Molecular detection of *B. microti* DNA in blood and tissue of females, dams, and their offspring

#### The frequency of B. microti DNA detection in female organs and blood

The comparison and statistical evaluation of differences in the frequency of *B. microti* DNA detection in blood and other tissues of females are presented in Table 4. DNA of *B. microti* was found in blood samples collected from all infected females in experimental groups and from Control Group NP (infected, non pregnant). *B. microti* DNA was identified in all tested organs (Table 4). The highest percentage of *B. microti* infected tissues was detected in hearts and lungs, with the lowest in uterus.

**Table 4.** Frequency of detection of *B. microti* DNA [Cl_95_] in blood and organs of BALB/c mice females from experimental groups and from the control group of infected, nonpregnant females.

| Speciment tested/sample | A<br>(n =6) | B<br>(n =15) | C<br>(n =20) | D<br>(n =20) | Control<br>Group NP<br>(n =6) | Statistical<br>values |
| --- | --- | --- | --- | --- | --- | --- |
| Blood | 100% [59-100] | 100% [27-94] | 100% [83-100%) | 100% [83-100] | 100% [59-100] | NS |
| Brain | 67% [27-94] | 53% [29-78] | 70% [48-86] | 100% [83-100] | 100% [59-100] | $\chi^2= 3.97^*$ |
| Heart | 100% [59-100] | 53% [29-78] | 70% [48-86] | 100% [83-100] | 100% [59-100] | $\chi^2= 4.97^*$ |
| Lungs | 67% [27-94] | 60% [33-81] | 85% [63-96] | 100% [83-100] | 100% [59-100] | $\chi^2= 3.54^*$ |
| Liver | 67% [27-94] | 27% [10-53] | 35% [17-58] | 90% [68-98] | 100% [59-100] | $\chi^2 = 8.29^{**}$ |
| Spleen | 67% [27-94] | 40% [19-67] | 60% [37-79] | 100% [83-100] | 100% [59-100] | $\chi^2 = 6.18^{**}$ |
| Kidneys | 67% [27-94] | 47% [22-71] | 55% [32-76] | 90% [68-98] | 100% [59-100] | $\chi^2 = 14.43^*$ |
| Uterus | 0% [0-41] | 13% [2-40] | 25% [10-48] | 70% [48-86] | 100% [68-100] | $\chi^2 = 10.30^{**}$ |

\* $p<0.05$ , $df= 4$ , \*\* $p<0.001$ , $df= 4$

#### The frequency of *B. microti* DNA detection in offspring organs, blood, and placentas

Organs were isolated from selected pups in Groups B (*n* = 14) and D (*n* = 33) on the 1^st^ day postpartum, and only from pups in Group B on the 7^th^ (*n* = 8) and 14^th^ (*n* = 19) day postpartum. From embryos, organs were isolated on the 12^th^ (*n* = 12 from Group B), 14^th^ (*n* = 8 from Group B and *n* = 12 from Group D), 16^th^ (*n* = 22 and *n* = 36, respectively), and 18^th^ (*n* = 29 and *n* = 21, respectively) day of pregnancy.

DNA of *Babesia* was found in all types of tested tissues (Table 3). Hearts, lungs, spleen, and placenta presented the highest *B. microti* positive rate. Higher positive rates in tissues of the liver, spleen, kidneys, and placentas was noted in offspring from Group D in comparison to Group B (Table 3).

#### Histopathological changes in female BALB/c mice

The material for histopathological examination was collected from selected females in experimental and control groups. A variety of histopathological changes (lesions) were found in organs of females in experimental and control groups (Table 5). There was no significant effect of the *Babesia* infection on the presence/frequency of lesions in female organs. Some of the detected changes were associated with aging or pregnancy development (results not presented).

**Table 5.**
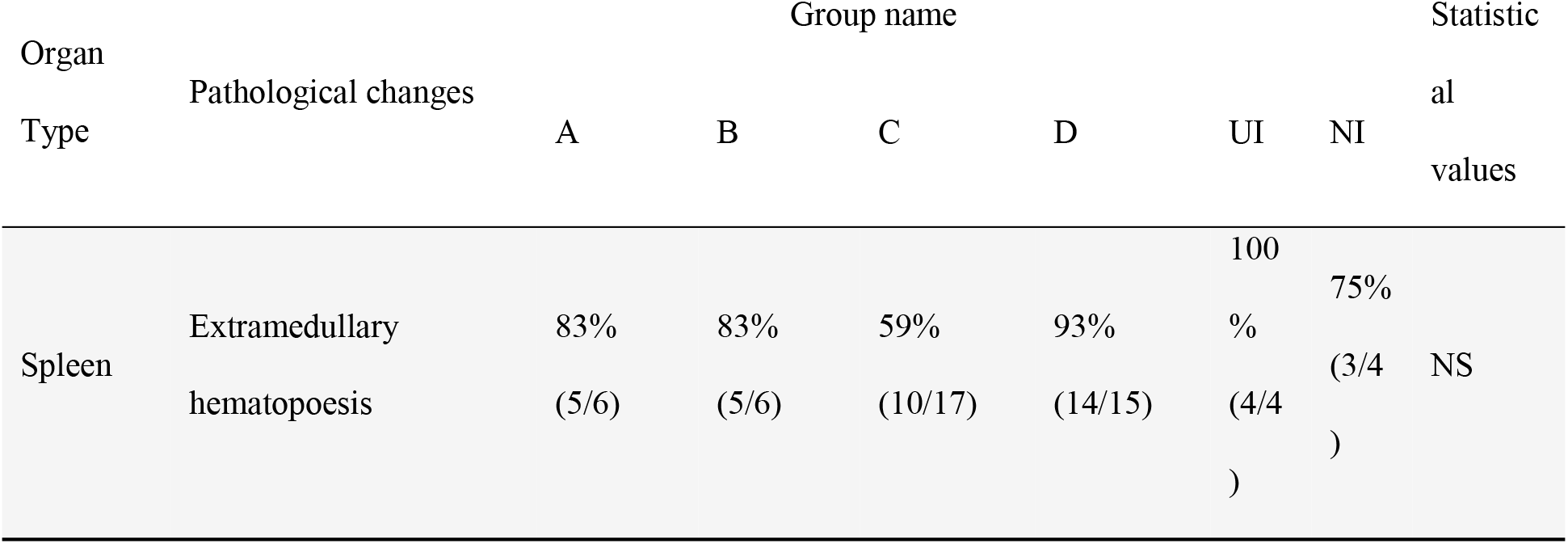
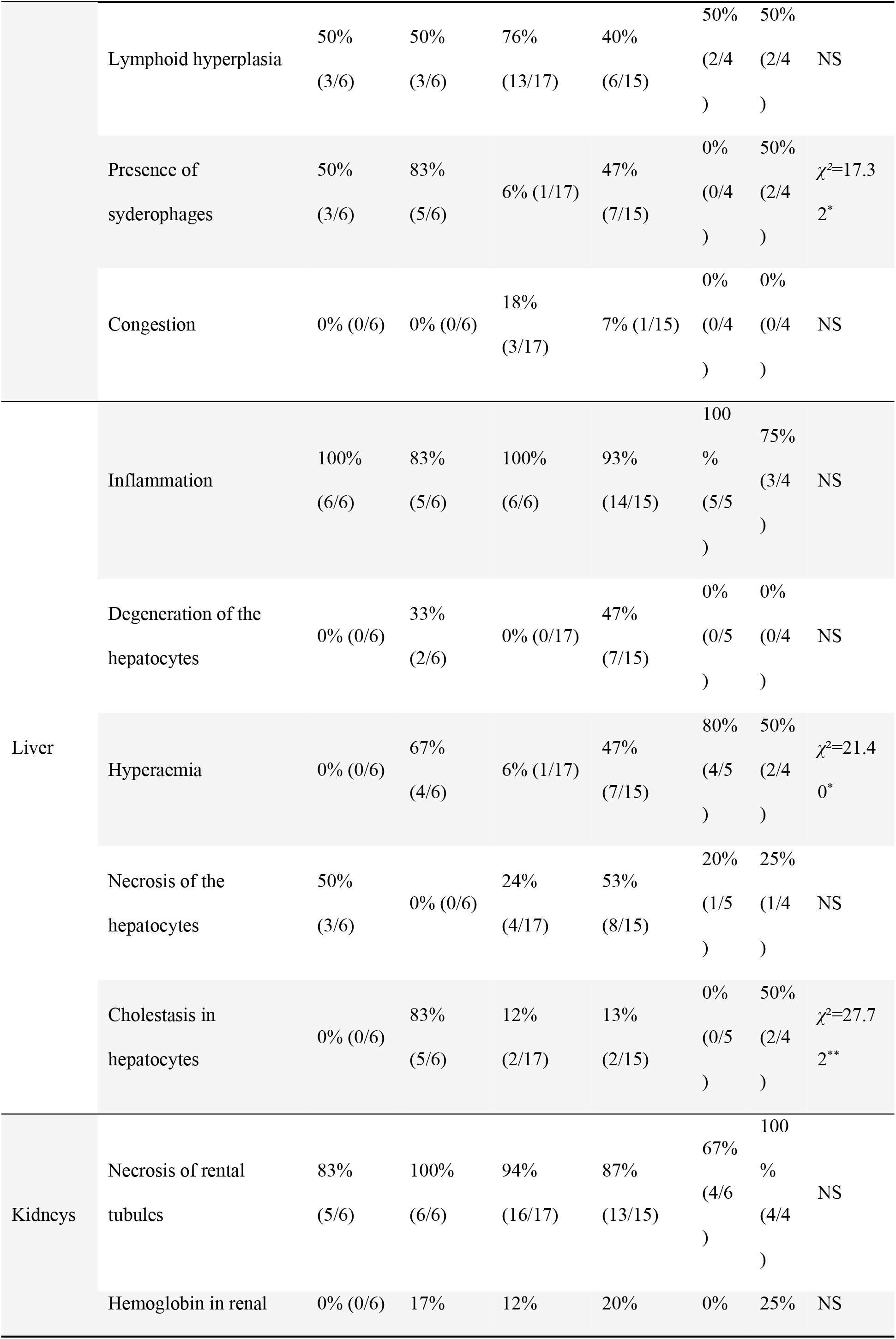
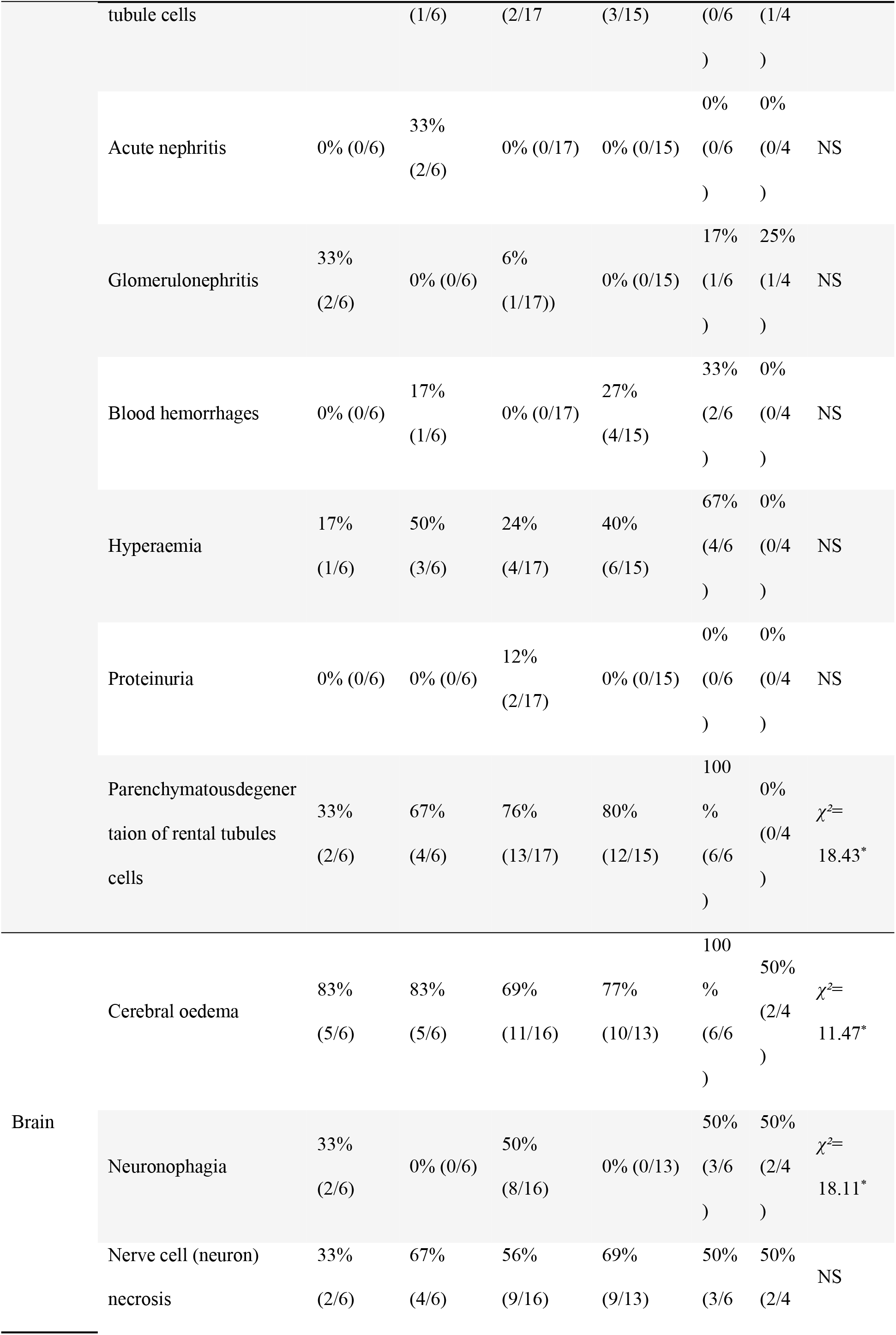
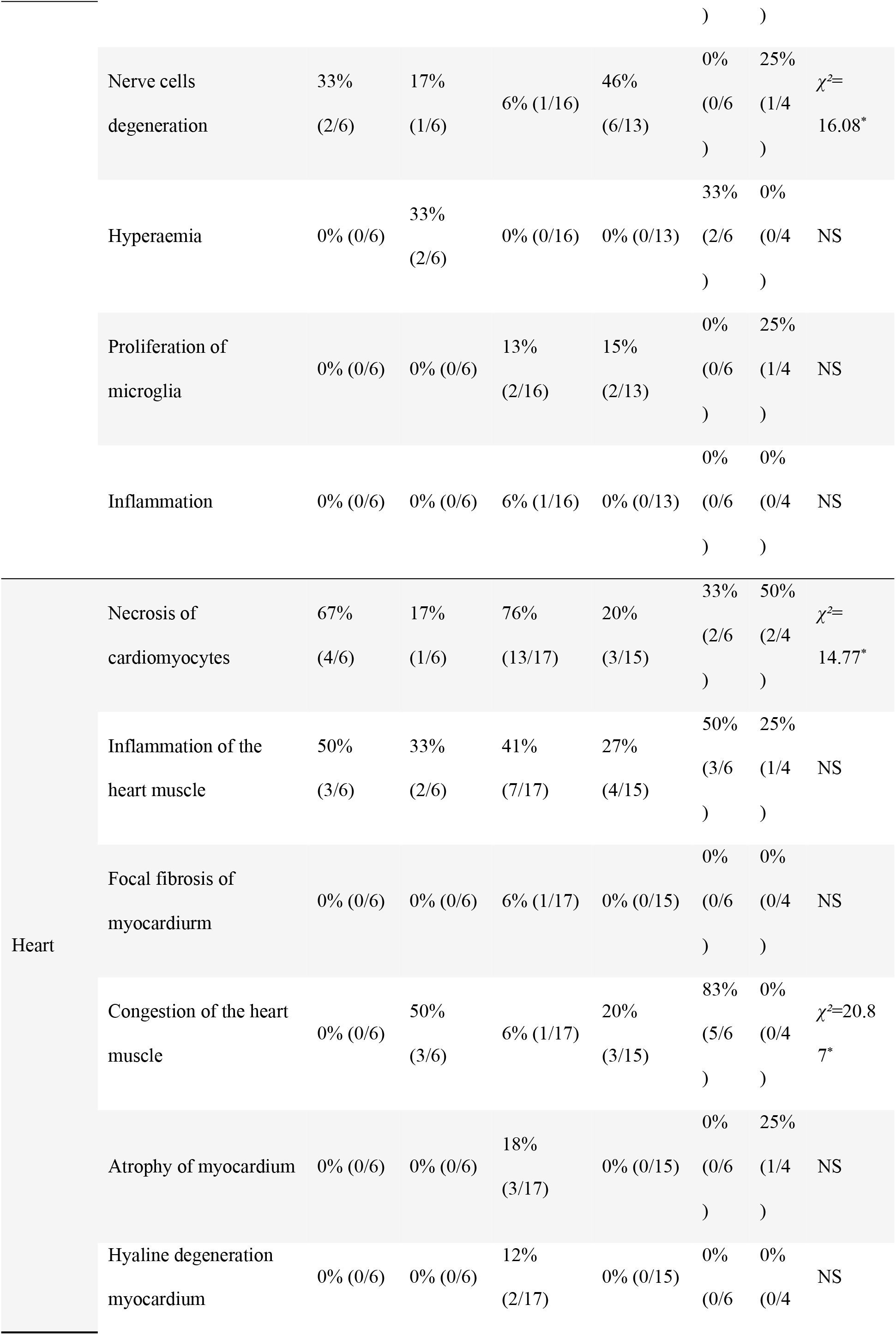
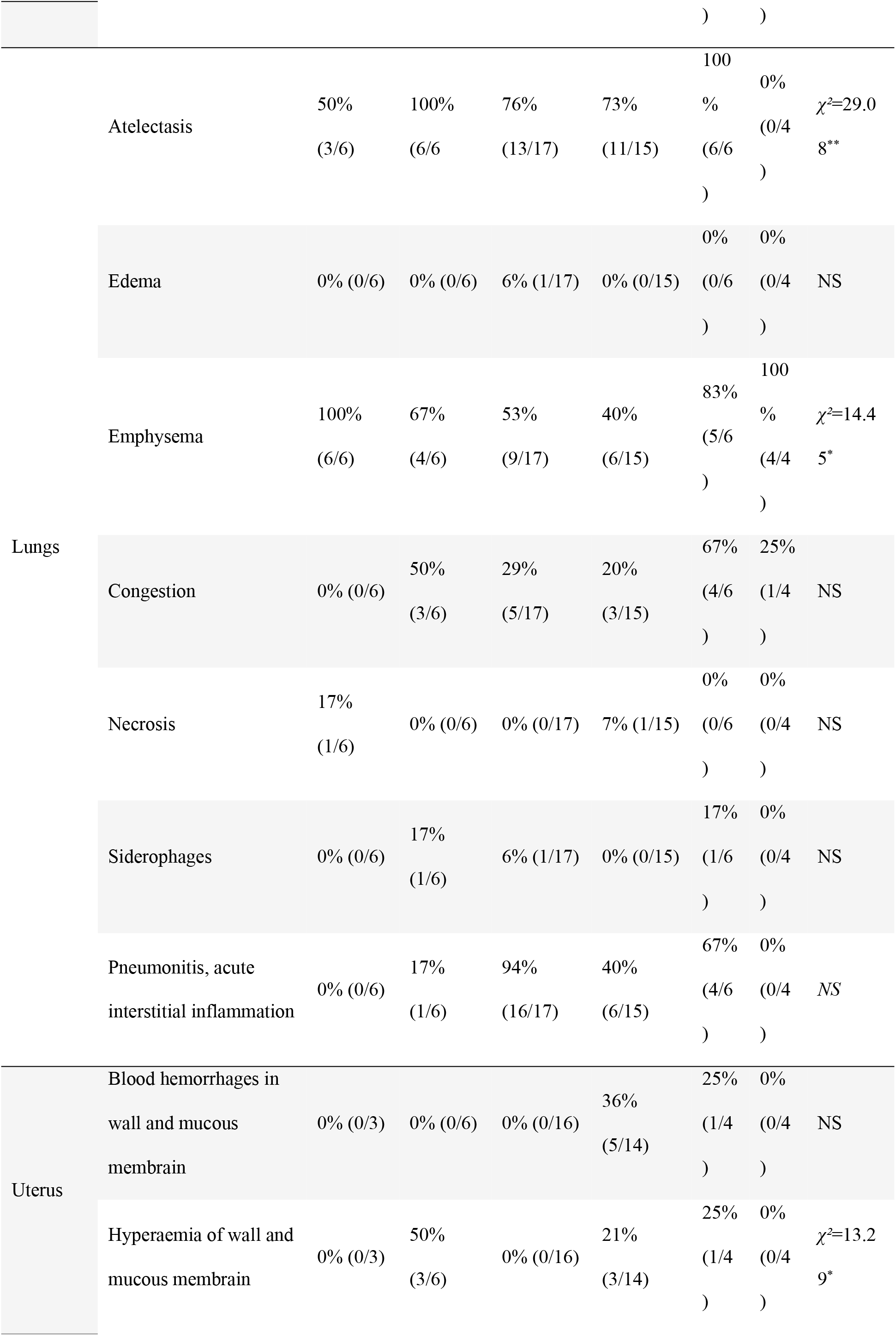
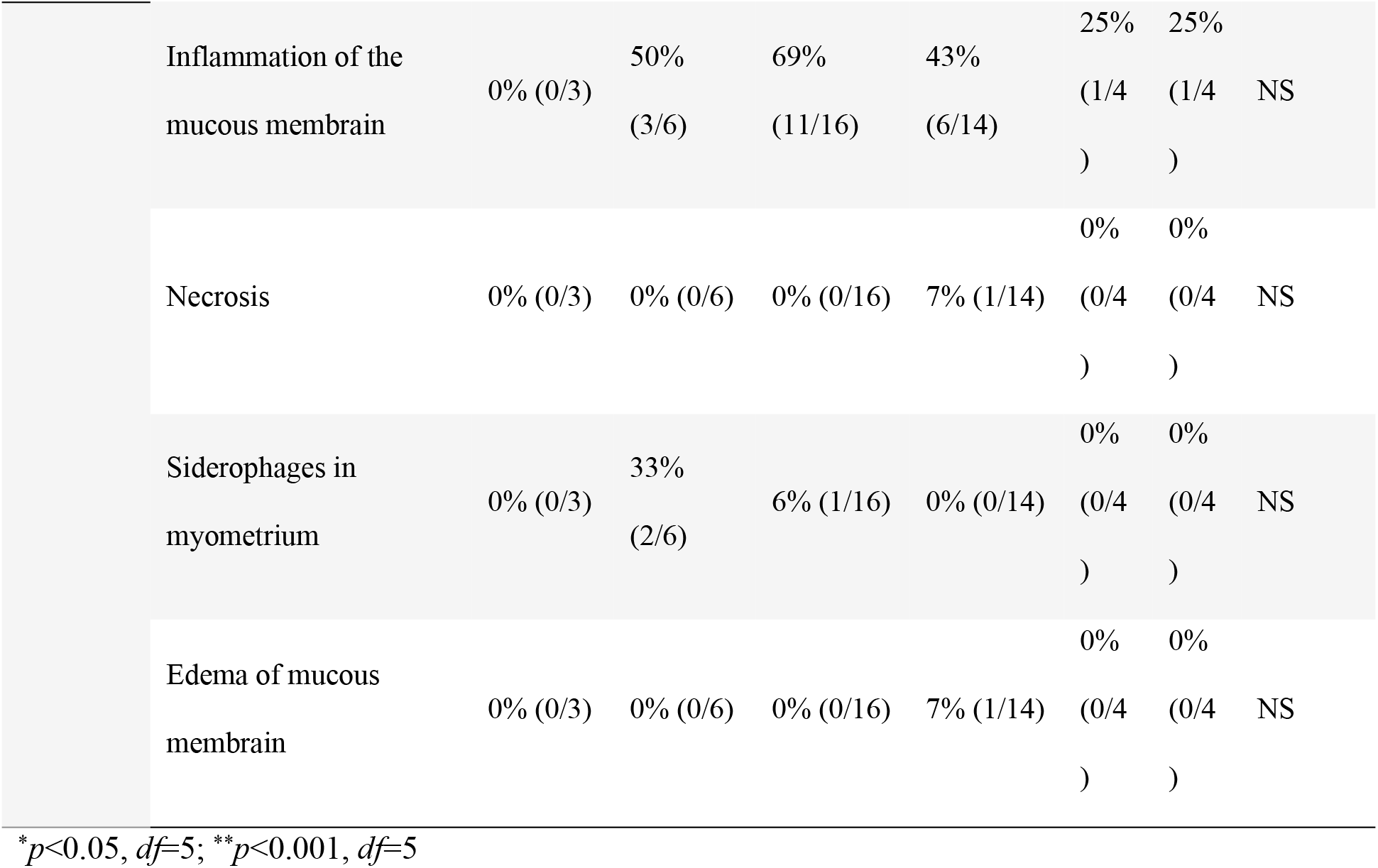
Mean number of pathological changes in tested organ tissues of females from experimental groups.

#### Histopathological changes in offspring

The material for histopathological examination was collected from selected pups in experimental Group B (*n* = 18) and Control Group UI (*n* = 12) on the 14^th^ day postpartum. A variety of histopathological changes were found in organs of pups in experimental Group B as well as in pups in Control Group UI (Table 6). There was no significant effect of the *Babesia* infection on the frequency/occurrence of lesions in organs of pups. Atrophy of cardiac myocytes occurred more frequently in heart tissues in which *Babesia* DNA was detected in comparison to negative samples (4/6=67% vs 1/11=9%; *χ²*=6.75, *df* = 1, *p*<0.05), but no other statistical association was found between infection (molecular detection) and pathological results.

**Table 6.**
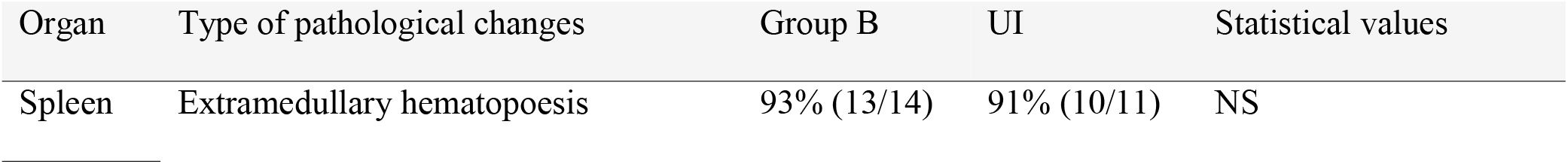
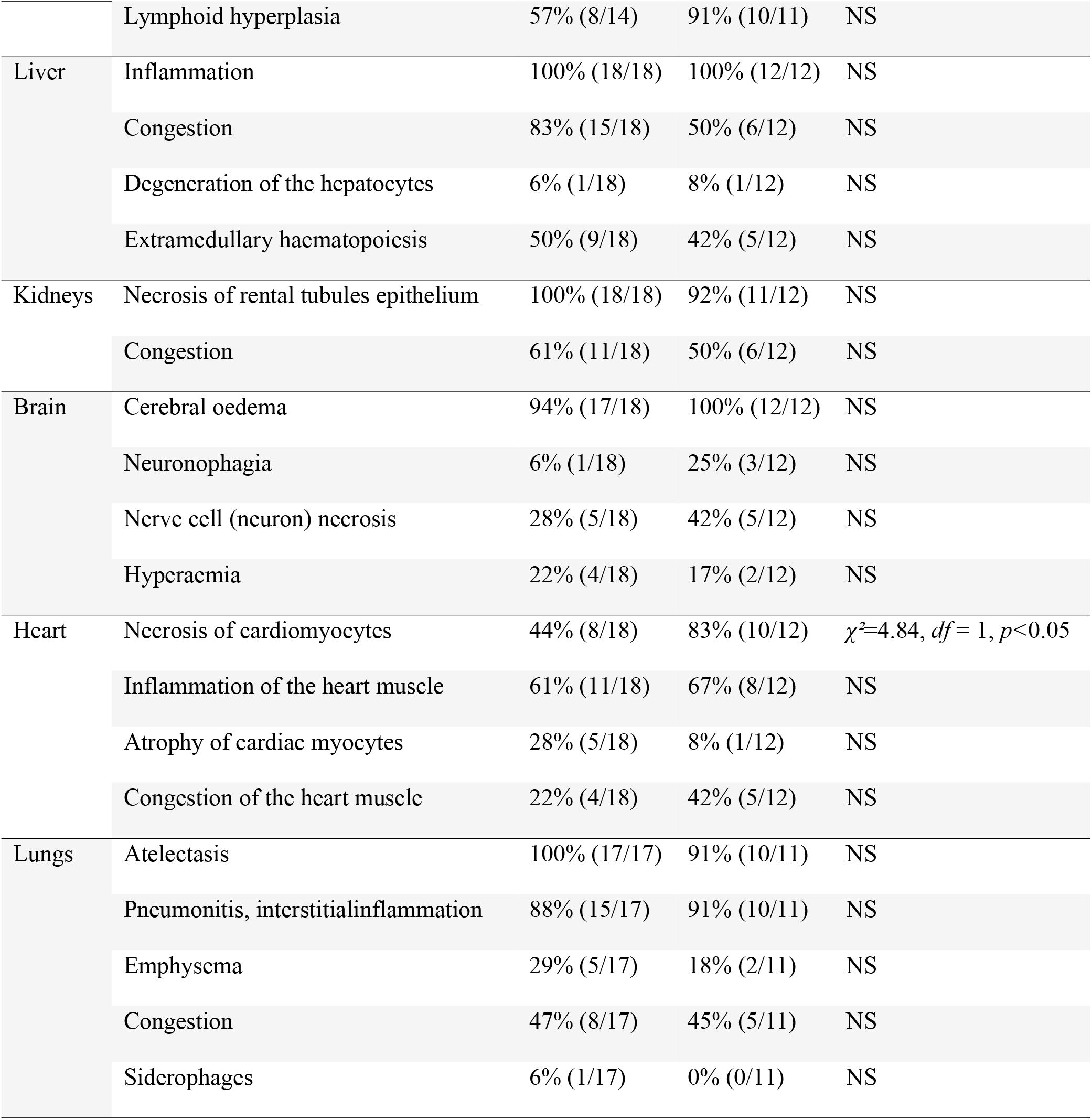
Frequency of pathological change occurrence in tissues collected from pups from Group B and Control Group UI on 14 day postpartum.

#### Blood parameters

We present the comparison of mean blood parameters for 6 females from each group (Fig 4). As females from Groups A and C presented no evidence of pregnancy, mean counts of RBC and platelets from these groups were compared to the counts obtained from Group NPI (non-pregnant, uninfected females) (Fig 4ab). RBC count was the lowest in Group A, and slightly higher in Group C. In both experimental groups, mean RBC counts were significantly lower than in the Control Group NPI (*t*=−17.92, *df*=10, *p*<0.001; *t*= −17.36, *df*=10, *p*<0.001, respectively). Platelet count was the lowest in Group C, and slightly higher in Group A. In both groups, platelet counts were significantly lower in comparison to the Control Group NPI (*t=* −3.69, *df*=10, *p*<0.05; *t*= −3.36, *df*=10, *p*<0.05, respectively).

**Fig 4.**
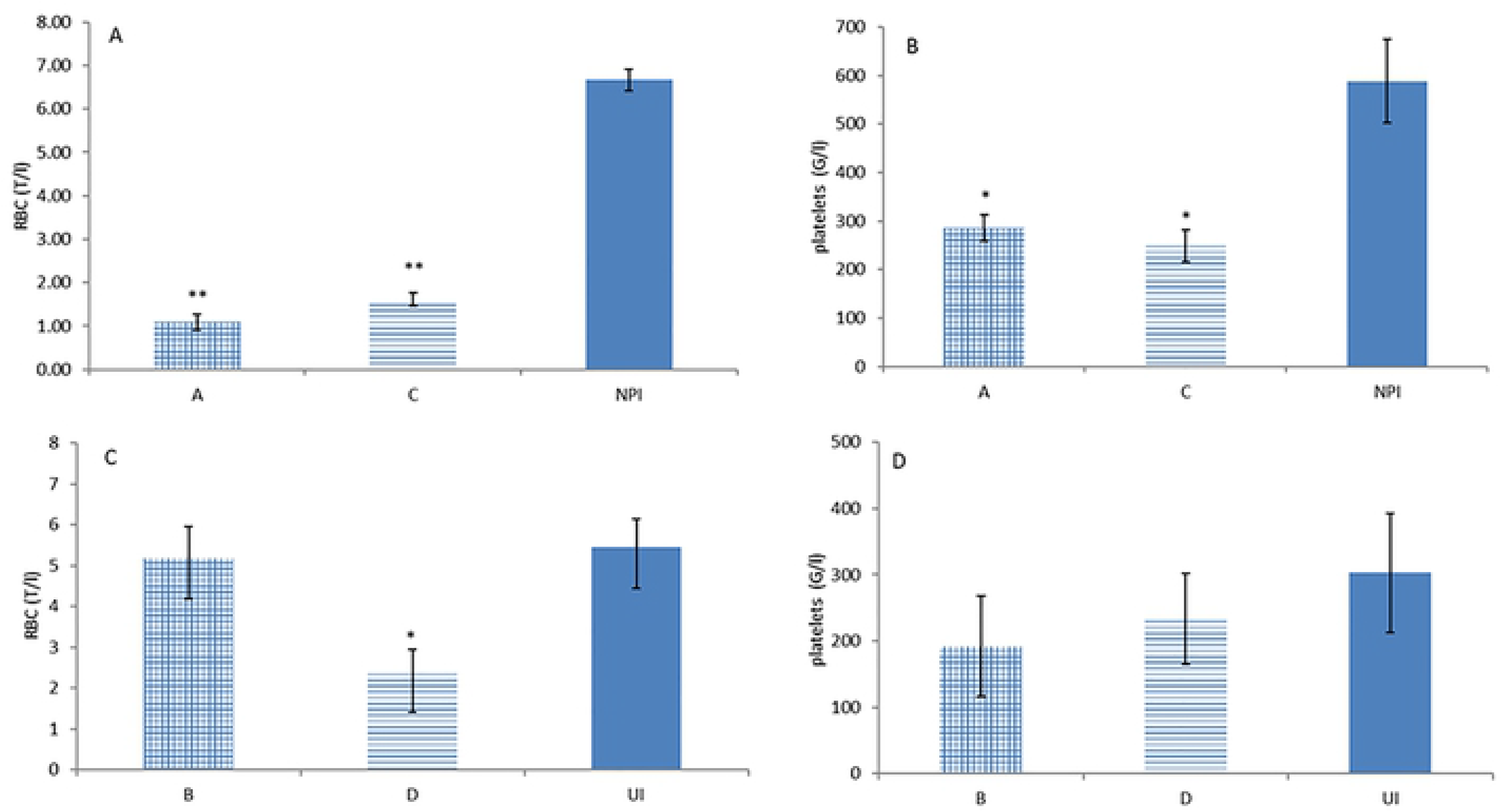
Comparison of the blood count results. A) RBCs and B) platelets in females mated on the 7^th^ day post infection (Group A), infected on the 4^th^ day post mating (Group C), in control group of infected, non-pregnant, uninfected females (NPI); C) RBCs and D) platelets in females mated after the 40^th^ day post infection (Group B), infected on the 12^th^ day post mating (Group D), and in control group of pregnant, uninfected females (UI).\**p*<0.05, ** *p*<0.001

Mean blood counts of females in Groups B and D were compared to the counts from Group UI (pregnant, uninfected females) (Fig 4cd). The mean RBC count (but not the platelet count) was higher in Group B in comparison to Group D (*t*=0.013, *df*=10, *p*<0.05). RBC count in Group D was significantly lower than in the Control Group UI– mean RBC 2.34±0.55 T/l and 5.44±0.69 T/l, respectively (*t*=−3.51, *df*=10, *p*<0.05).

## Discussion

In this study, we have confirmed a significant negative influence of *B. microti* infection on the initiation of pregnancy in females in the acute phase of infection, but a lack of negative influence of chronic infection on pregnancy development. In our study, *B. microti* infection did not show significant influence on histopathological condition of tested tissues.

In females infected shortly before embryo implantation (Groups A and C), pregnancies did not develop or were resorbed (presence of scars) during the first trimester. Negative impacts of the infection on the initiation of pregnancy might be the result of high parasitaemia level and significant decrease in maternal RBCs and platelets in experimental Groups A and C, resulting in severe anemia and thrombocytopenia. Results presented in this study support the outcomes of previous work, where females mated on the day of infection with *B. microti* did not deliver pups [17].

Our experiments confirmed the negative influence of acute *Babesia* infection on pregnancy development and the reproductive success of mice. Females from Group D were infected in the second trimester and delivered infected pups, which were unable to survive longer than 1-2 days post-delivery. Mortality was reported to occur between the 1^st^ and 30^th^ day after birth in infants untreated for congenital toxoplasmosis, Chagas disease, and malaria [3, 35, 36]. *Plasmodium* infection in female mice caused the low birth weight of pups, the increased frequency of abortion, and the greatly increased foetal death rates observed in murine malaria model [37]. Similarly, in non-immunised pregnant women, malarial infection during the first or second trimester has been associated with high rates of abortion [38]. *Plasmodium vivax* infections are associated with an increased chance of miscarriage, the occurrence of intrauterine growth restriction [39, 40], and lowered birth weight [41, 42]. Analysis of the relationship between the time of mating and the phase of infection revealed that the pups were delivered only by females with post-acute or chronic infections [17]. In this study, the reproductive success of chronically infected females was much higher than in our previous study [17]. This result may be the outcome of different approaches in study design. Extended cytological examination in this study enabled us to increase reproductive success by mating females exactly in the oestrus stage. In the previous study, female reproductive status was judged by visual assessment of the vaginal opening, then males were joined with females for a few consecutive nights for mating [17]. These changes in the study procedures may have increased the number of pregnant females in the current study. In the group of mice with chronic babesiosis infection (Group B) we did not record abnormalities in the reproductive success or in offspring condition. Pups delivered in Group B presented no symptoms of worsened condition (their heartbeat rate during pregnancy and body mass postpartum were normal).

An interesting observation in our study is that the probability of successful vertical transmission increased with the duration of the pregnancy. In females infected in the second trimester, *B. microti* transmission was recorded since the 14^th^ day of pregnancy (2^nd^ day post infection). Our results suggest that the success of vertical transmission is the highest in the third trimester, when placental tears may favor parasitic transmission [43]. The previous study revealed that the success of vertical transmission was dependent on the phase of the parasite infection [17]. Since the circulation between the female and the fetus is established only on day 9 or 10 of gestation in mice, vertical transmission of parasites is expected to occur rarely in the first trimester [19, 44]. It is possible that placental breaches/tears resulting from damage induced by placental inflammatory responses or appearing naturally close to the pregnancy termination, particularly during labour, can facilitate congenital transmission of parasites from maternal blood [19]

The success of vertical transmission of *B. microti* estimated in this study was markedly lower than in our previous work [17]. It might be linked to the time of blood collection and changes in maternal and individual immunity in pups. In the current experiment, blood samples were taken between the 12^th^ day of pregnancy and 14^th^ day postpartum. In our pervious experiments, blood samples were taken after the 28^th^ day postpartum [17]. Interestingly, infants with congenital babesiosis develop symptoms and are admitted to hospital following 4-6 weeks postpartum (fever, fatigue, irritability, and decreased oral intake) [24, 45]. A similar delay in appearance of babesiosis symptoms was observed in pups with congenital infection of *B. canis* – at 6 weeks, a period of life that is believed to be crucial in the development of immunity in dogs [13]. During this period, maternally derived immunity decreases rapidly, to enable the development of individual immunity [46]. It is possible that in the mouse model, decreasing derived maternal immune response gives *B. microti* chance to multiply if the individual immunity is not strong enough.

The course of parasitaemia in all experimental groups was typical for *B. microti* infection with a decline after the 30^th^-40^th^ day post infection. However, molecular examination still indicated an ongoing infection. These results are in accordance with the hypothesis proposed by Bednarska et al.2015 [17] that states that parasites can modulate the immune system and when chronic infection is established and well stabilised in mice, the hormonal changes associated with the development of pregnancy do not affect the course of the *B. microti* infection. In our model, pregnancy and lactation did not significantly change the level of parasitaemia in females in the chronic phase of infection. It was also observed that several females that were pregnant and lactating in the post-acute phase of *B. microti* infection showed minor increases of parasitaemia, but it was not recorded in this study [17].

In protozoan infections, parasitaemia levels in pregnant women/females seems to be an important factor contributing to occurrence of vertical transmission, e.g. congenital infections with *T. gondii*, *T. cruzi, P. falciparum* occur more frequently in mothers displaying higher parasitaemia [35, 47–49]. However, we did not find any significant influence of parasitaemia level on vertical or reproductive success of mice.

DNA of *B. microti* was detected in all types of tested organ tissues of females and their offspring. The highest percentage of *B. microti* infected tissues was detected in the hearts and lungs, with the lowest in uterus. DNA of *B. microti* was detected in all tested types of tissues collected from pups and embryos from Groups B and D. The highest percentage of *Babesia*-infected tissues was detected in hearts, spleens, and placentas. Sequestration of iRBCs in spleen, lungs, and hearts was described in humans infected with *B. microti,* as well as in bovines infected with *B. bovis,* and hamsters experimentally infected with *Babesia* WA1 [50–52]. Poovassery and Moore (2006) described the accumulation of *Plasmodium chabaudi* AS-infected erythrocytes in the placenta of infected mice as a manifestation of specific placental sequestration [53]. It is possible that placental accumulation is a phenomenon also observed in *Babesia*-infected placental tissues. High parasitaemia in placental blood, while lower in newborns, was observed in malaria congenital infection [3]. The reported incidences of congenital malaria in endemic areas of Africa and Asia are highly variable. Most reported rates of maternal–fetal malaria transmission in range 30% - 55% were obtained from detection of parasites in cord blood ([3, 54–57]. Much lower rates, up to 0.5% ([58], were reported from studies on infant peripheral blood, suggesting a rapid elimination of congenitally-transmitted iRBC. The possible reasons of such self-cure of malaria congenital infection are still unknown. Most newborns with congenital malaria are able to control and clear the infection without receiving specific treatment [19], however, mechanisms leading to such favourable self-cure are poorly recognised. They may involve neonatal immune responses toward parasite, as well as maternally transmitted antibodies and the presence of high levels of fetal haemoglobin [3, 54, 59]. Further investigation considering maternal and neonatal immune response is necessary to find out the immune response course in case of congenital babesiosis.

*B. microti* infection causes the decrease in RBCs counts because of both the sequestration of infected cells and disintegration of the iRBCs during multiplication of parasites [60]. For these reasons, anemia is one of the main symptoms of babesiosis and it was observed in females from Groups A, C, and D in the acute phase of infection. In addition to the immune response and placental accumulation of parasites, advanced anemia may play a role in the observed pregnancy loss in our experiments (i.e. because of limited oxygen transportation to developing fetuses). As a result of the self-healing process, the number of RBCs increases shortly after the acute phase, and its normal during the chronic phase of infection [7], which was observed in Group B. Development of parasitaemia and anemia were accelerated in *P. chabaudi* AS-infected pregnant mice [53]. Acute babesiosis in women leads to severe anemia and thrombocytopenia if diagnosed in second or third trimester [61, 62].

In conclusion, acute *B. microti* infection prevents pregnancy initiation and/or development of pregnancy at a very early stage, and causes severe complication in BALB/c mice in the second and third trimester of pregnancy. Chronic *B. microti* infection has no negative impact on the initiation and development of pregnancy, but resulted with congenital infections. Further study is required to determine to what extent maternal antibabesial immune responses and potential placental accumulation of parasites contribute to compromised pregnancy in the murine model of congenital *Babesia* infection.

## Acknowledgments

We thank our colleagues and students who provided insight and expertise that greatly assisted the research. We thank: Joanna Zajkowska, Dr Katarzyna Szczepańska, Dr Aneta Suwińska and Dr Katarzyna Kisiel. We thank Professor Francis S. Gilbert from the University of Nottingham, UK for sharing the statistical software. Special thanks to Stephen J. A. Jennings for proofreading the article.

